# Impaired thalamic burst firing in Fragile X syndrome

**DOI:** 10.1101/2025.01.27.635156

**Authors:** Ronan T. O’Shea, Nicholas J. Priebe, Darrin H. Brager

## Abstract

The thalamus performs a critical role in sensory processing by gating the flow of sensory information to the neocortex and directing sensory-driven behaviors; functions which are disrupted in people with autism spectrum disorders (ASD). We have identified cellular changes in thalamic neurons in a mouse model of Fragile X syndrome (FX), the leading monogenic cause of ASD, that alter how the thalamus transmits sensory information to neocortical circuits. In awake animals, thalamic relay cells gate input by shifting between two firing modes: burst and tonic. Relay cells in FX mice, however, do not shift between these modes and instead operate primarily in the tonic mode. We demonstrate that the lack of burst mode firing is caused by a shift in the voltage sensitivity for the Ca^2+^-dependent low threshold spike, which underlies normal burst firing.

## Introduction

In the face of complex sensory environments, animals must segment task-relevant features from the background to guide actions and avoid becoming overwhelmed by irrelevant noise. What constitutes target and background in the sensory landscape can vary rapidly, making flexible control over what sensory information gets processed by neocortical circuits crucial for guiding optimal behaviors.

People with ASD commonly struggle to detect, attend to, and interpret salient stimuli. This impairment is due to feeling overwhelmed by uninformative stimuli in “noisy” environments, hypersensitivity to normally innocuous features, and trouble redirecting attention when salient stimuli appear. Psychophysical studies demonstrated that these sensory abnormalities cannot be explained by changes in detection thresholds for low-level features^1,2^. Rather, there is a deficit in higher-order processing, apparent in tasks requiring the segmentation of task-relevant features from background ^2–7^. This sort of hyposensitivity to behaviorally relevant stimuli in ASD stands in contrast with widely reported perceptual hypersensitivity to supra-threshold stimuli, which correlates with larger magnitudes of neocortical activation than control groups, as measured with EEG ^8–10^. These reports of both hypo- and hypersensitivities suggest that the relay of sensory information to neocortex is dysregulated in ASD.

Given these sensory abnormalities, it is possible that typical relay of sensory information from the periphery to neocortex, via the thalamus, is disrupted in ASD. While the thalamus is often considered a relay station, it also acts as a critical gate of information: the statistics of sensory inputs and modulatory feedback from neocortex and brainstem alter how the thalamus conveys sensory information to the neocortex^11–13^. These inputs alter information transmission by shifting thalamic relay cells between two modes of action potential (AP) firing: burst and tonic modes^14^. Bursting is mediated by the generation of the low threshold spike (LTS), a non-linear depolarization of membrane potential mediated by T-type Ca^2+^ channels, which are activated at membrane potentials near rest^15–18^. The combination of the statistics of the sensory input and top-down modulatory drive sets the membrane potential of a thalamic relay cell, which determines whether it will fire in the burst or tonic mode in response to a stimulus^19–21^. Burst and tonic firing play complementary roles in relaying sensory information^13,22–26^.

The proposed role for thalamic firing dynamics in selectively gating sensory relay to neocortex and the behavioral evidence for dysregulated sensory processing in ASD led us to compare thalamic population activity in wild type (WT) and FMR1-knockout (FX) mice, a common experimental model for exploring the physiological basis of the autistic phenotype^27^. We found that burst firing *in vivo* is profoundly impaired in the FX thalamus, with these relay cells firing almost exclusively in the tonic mode, in stark contrast with the distribution of firing modes which we recorded in WT thalamus. To investigate the basis for impaired burst firing in FX at the cellular level, we performed *in vitro* whole-cell recordings from relay cells in acute brain slices and found that the voltage sensitivity of the LTS was hyperpolarized in FX thalamic neurons, which contributes to reduced burst frequency in FX thalamus.

## Results

To compare the thalamic firing patterns in WT and FX mice, we densely sampled single unit activity in LGN of awake WT and FX mice with the Neuropixels 1.0 probe^28^. We classify spikes as emerging from burst or tonic firing mechanisms by the preceding and subsequent ISI: the first AP in a burst is preceded by at least 100ms of silence, followed by APs separated by less than 4ms; ending with an ISI longer than 4ms; all other spikes are classified as tonic (Fig. 1A)^29^. The pattern of tonic and burst spiking is consistent across WT relay cells (Fig. 1B). In contrast, relay cells in FX LGN fired primarily in the tonic mode (Fig. 1C). The frequency of burst events was significantly lower in FX than WT populations during both spontaneous and visually-evoked activity (WT spontaneous mean=.16 +/- .25, median=.07 bursts/sec; FX spontaneous mean=.01+/- .03, median=.004 bursts/sec; p<.01 for a Wilcoxon rank sum test; WT evoked mean=.09+/- .11, median=.04 bursts/sec; FX evoked mean=.01+/- .03, median=.004 bursts/sec; p<.01 for a Wilcoxon rank sum test) (Fig. 1D, left). Even when FX relay cells did burst, the bursts contained significantly fewer APs compared to bursts from WT relay cells (WT mean= 2.94 +/-1.02 spikes/burst, FX mean= 2.46 +/-.81 spikes/burst; p<.01 for a Wilcoxon rank sum test) (Fig 1F).

**Figure 1:**
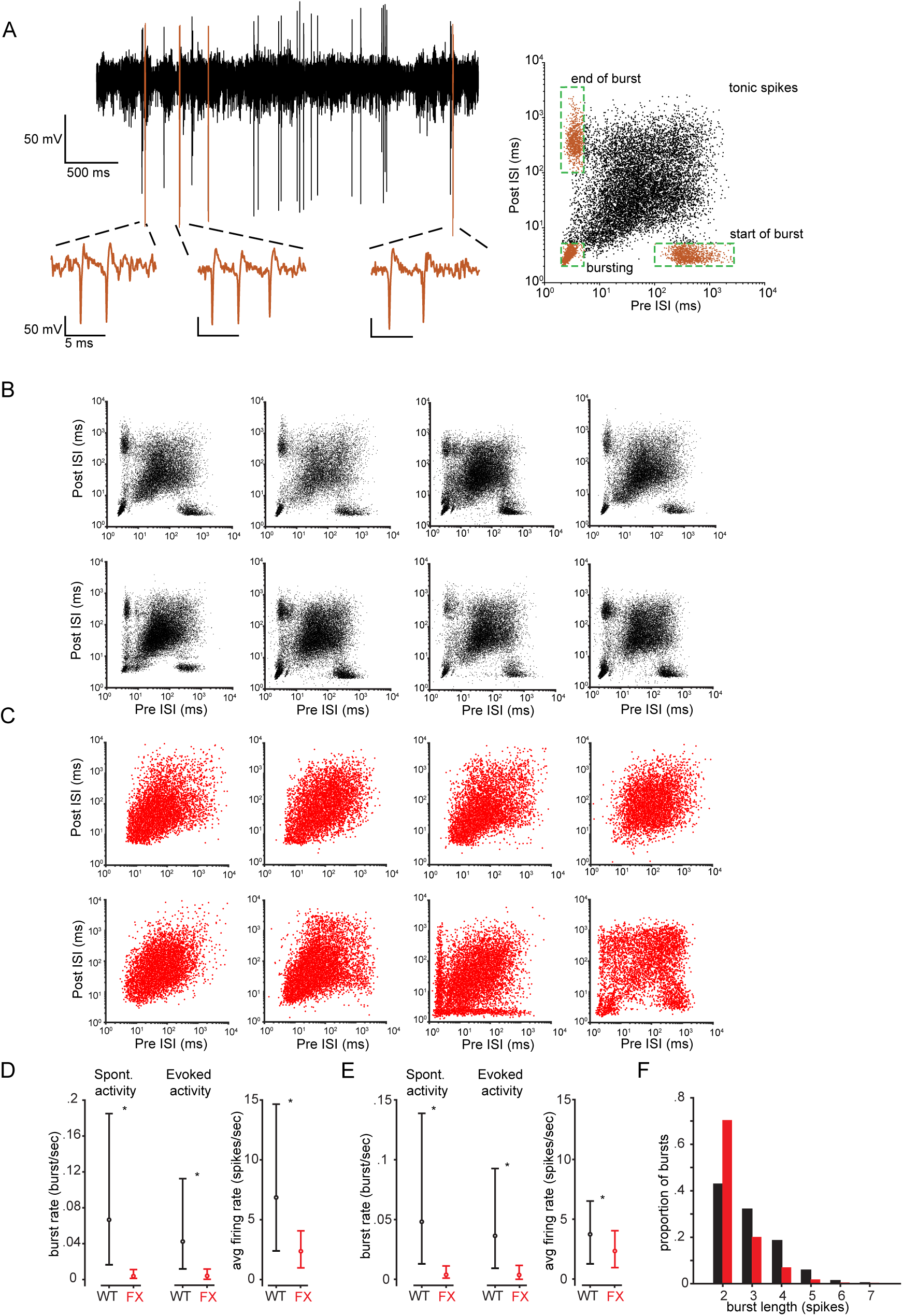
Firing patterns in LGN of awake WT and FX mice. **a**. (left) Example voltage trace of WT LGN relay cell exhibiting tonic (black) and burst (orange) spikes. (Right) Example ISI distribution of WT LGN relay cell, where each scatter point represents the pre and post ISI for a single spike. Green boxes indicate the criteria for spikes being classified as part of a burst. **b**. Example ISI distributions of WT LGN relay cells. **c.** Example ISI distributions of FX LGN relay cells. **d**. (Left) Distribution of burst rates for WT (black) and FX (red) populations. (Right) Distributions of baseline firing rates. Circles are the medians of the distributions. Lower and upper bars indicate the 25^th^ and 75^th^ percentiles of the distributions, respectively. *: p<.01 for a Wilcoxon rank sum test between FX and WT population data. **e**. As in **a**, for the resampled WT dataset. **f**. Histogram of the number of spikes in each burst across the WT and FX LGN populations.

WT relay cells also had higher baseline firing rates than FX relay cells (WT mean=10.3 +/- 10.4 spikes/sec, median=6.9; FX mean=3.2 +/-3.3, median=2.4 spikes/sec) (Fig. 1D, right). This difference in baseline firing rate could underlie the difference in burst frequency which we observe between the genotypes. To explore this possibility, we resampled our WT dataset to create a sample with firing rate statistics matched to the FX dataset (Fig. 1E, right). The distribution of burst frequencies for the resampled WT population remained significantly higher than that of the FX population, indicating that the disruption of thalamic firing dynamics in FX cannot be explained simply by a difference in the baseline firing rate of neurons across genotypes (WT resampled spontaneous mean=.13 +/- .19, median=.05 bursts/sec; WT resampled evoked mean=.07+/- .10, median=.04 bursts/sec) (Fig. 1E, left). These differences in thalamic firing dynamics between genotypes could also not be explained by differences in locomotion, as WT and FX mice spent equal amounts of time running during recordings.

The ISI distributions for FX relay cells show clear qualitative differences from those of WT relay cells. Though some FX relay cells do exhibit firing patterns which meet the criteria to be classified as bursts (Fig 1C, bottom right). To quantify thalamic firing dynamics of WT and FX populations we examined the interactions between successive spikes. Spike trains modeled by a Poisson process are characterized by an exponential distribution of ISIs. But neurons can deviate from this process in characteristic ways. After an AP there are refractory periods in which a successive AP is less likely, and bursting periods in which successive AP is more likely. One way to quantify the degree of burstiness and refractoriness in a neuron is to measure the deviation of the ISI distribution from an exponential fit^30^. While refractoriness results in relatively few spikes with short ISIs compared to the Poisson prediction, bursting generates spikes with short ISIs with an increased probability of spiking relative to the Poisson expectation for a given mean firing rate (Fig. 2A). We established a bursting weight for each relay cell by taking the ratio of the empirical ISI distribution to the rate-matched Poisson expectation within the 1-4 ms range (Fig. 2B). A weight of greater than 1 indicates that a cell bursts more frequently than expected for a Poisson process. A weight of less than 1 indicates that a relay cell fires primarily in the tonic mode, with the empirical ISI distribution showing relatively few spikes separated by small ISIs compared to the Poisson prediction due to the effects of the absolute and relative refractory periods.

**Figure 2:**
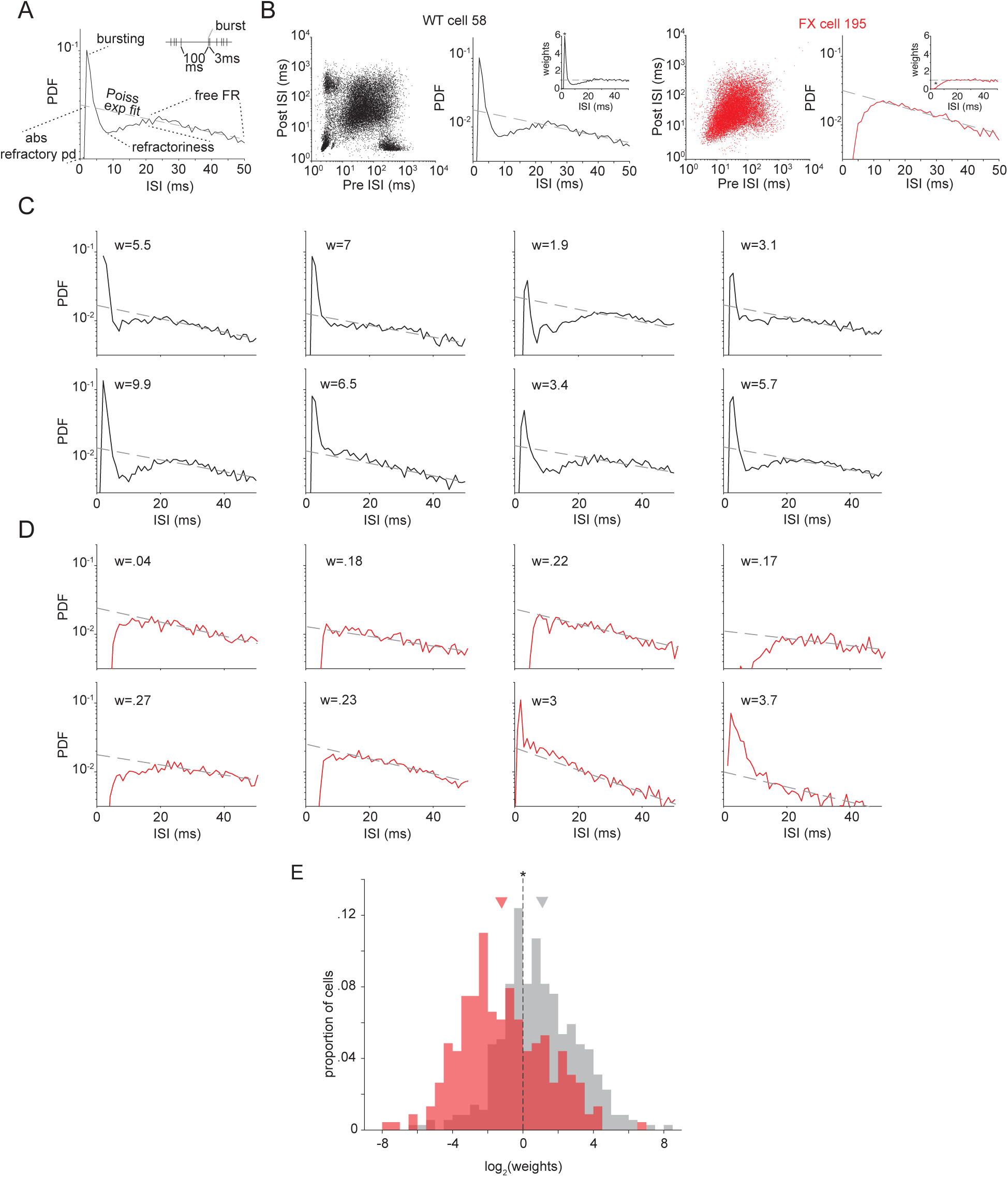
Deviations from Poisson statistics in WT and FX LGN populations: **a**. Example ISI PDF for a bursting relay cell. Black solid line is the empirical ISI PDF of the cell, gray dashed line is the ISI distribution of a Poisson process with a matched mean firing rate. **b**. Example ISI distributions for a WT (black) and FX (red) relay cell. (Inset) bursting weights computed for each example cell. **c**. Example ISI distributions for the same WT relay cells as in Fig 1B, with gray dashed line indicating the rate-matched Poisson expectation. Inset gives the bursting weight for each example cell. **d**. As in **c**, for the same example FX relay cells as Fig 1C. **e**. Histogram of log_2_ bursting weights for all relay cells in the WT and FX populations. Arrows indicate means of distributions.

This metric captures the stark difference in thalamic firing patterns between WT and FX populations. WT relay cells exhibit more spikes separated by 1-4ms than the Poisson prediction, indicated by bursting weights greater than 1 (WT geometric mean bursting weight = 1.87) (Fig. 2C,E). By contrast, burst spikes were infrequent in FX relay cells, indicated by bursting weights less than 1 (FX geometric mean bursting weight =0.47) (Fig. 2D-E).

One potential mechanistic explanation for impaired bursting in FX LGN is that the function of voltage gated T-type Ca^2+^ channels, which switch thalamic relay cells to burst mode at hyperpolarized membrane potentials, is altered in FX ^15,16,31–33^. To examine this possibility, we performed *in vitro* electrophysiological recordings from thalamic neurons in WT and FX mice. We made whole cell recordings from neurons in both the LGN and medial dorsal thalamus and all results reported were consistent across these two nuclei. Both WT and FX neurons showed an increase in AP firing rate in response to depolarizing current injections (Fig. 3a). WT neurons however, fired fewer APs compared to FX neurons particularly at the smaller current injections (Fig. 3b, mixed-factor ANOVA, interaction between genotype and amplitude: F (11,195) = 3.818, p < 0.0001). At larger current injections, firing rate increased as a linear function of current amplitude, in line with the observation from *in vivo* recordings that the tonic mode represents a linear relay of sensory inputs^14^. When neurons were held at -60 mV (near the resting V_m_) however, WT neurons reliably fired a burst of APs in response to small, brief current injections (Fig. 3c). In the burst mode, these neurons integrated inputs in a nonlinear manner, as increasing the duration of the current step up to 3000ms did not elicit more spikes after the initial burst, consistent with the proposed gating function of burst spiking. In contrast, FX neurons fired in the tonic mode and increased the number of spikes fired with increasing current duration (Fig. 3C, D; mixed-factor ANOVA, interaction between genotype and duration: F(5,80) = 8.193, p < 0.0001).

**Figure 3:**
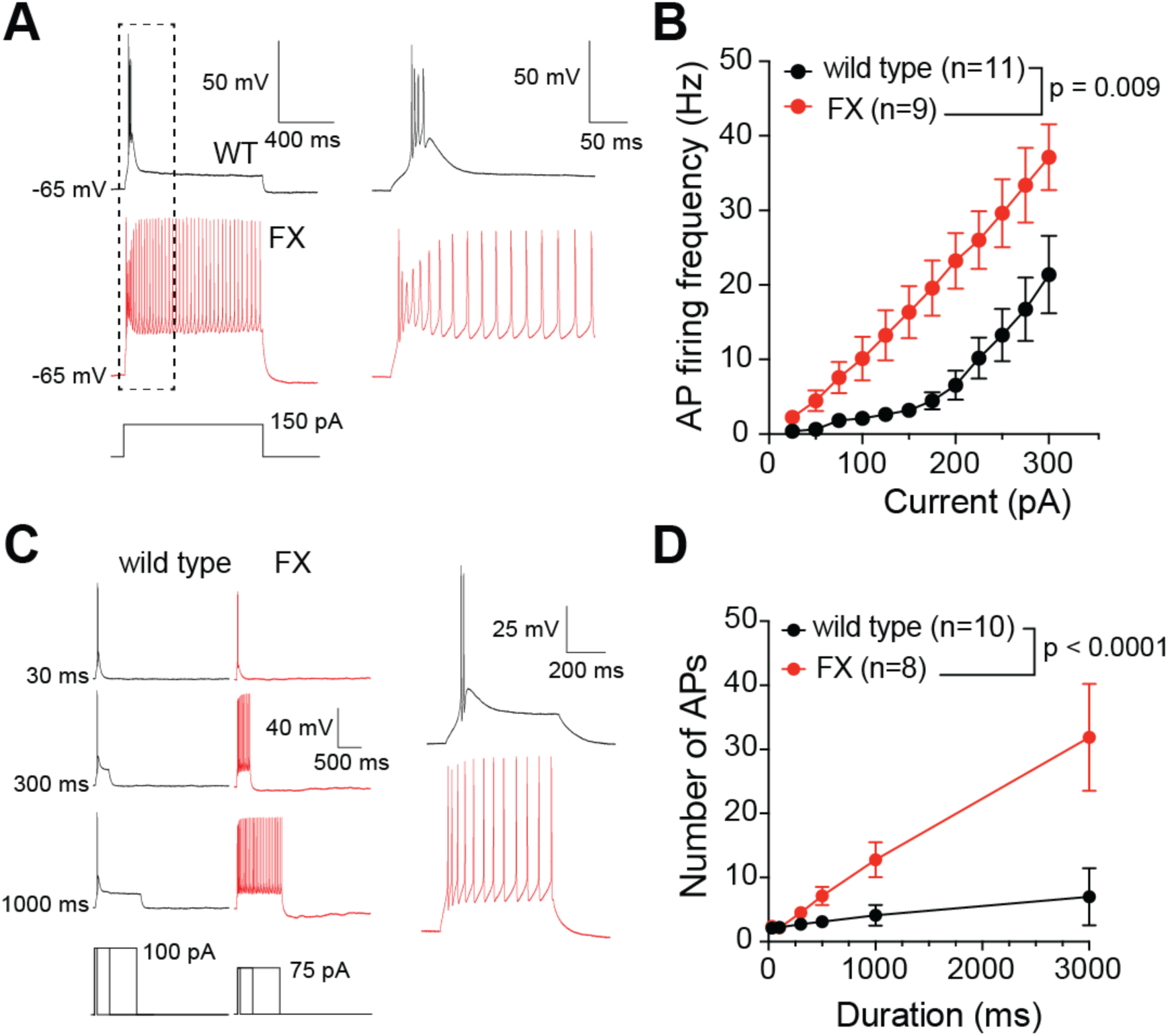
Thalamic neurons preferentially fire in tonic mode in FX. **A.** Action potential firing recorded from single thalamic neuron in response to current injection. Dashed box shows initial action potentials expanded at right. **B.** FX thalamic neurons fire more APs compared to WT. **C.** Action potential firing in response to current injections of variable duration. **D.** FX neurons show a linear increase in firing frequency consistent with tonic firing mode while WT thalamic neurons fire a brief train of 2-3 APs consistent with burst firing mode.

At rest, WT thalamic neurons respond to excitatory drive in the burst mode, amplifying sensory drive following a period of inactivity, whereas FX thalamic neurons respond to same excitatory drive in tonic mode^22,25^. There were no differences in the passive or AP properties between WT and FX neurons suggesting that the differences in burst firing were not likely due to Na^+^ or K^+^ channel dependent mechanisms (Supplementary Table 1).

The lack of bursts in FX relay cells at rest could be due to an absence of the LTS altogether, or a change in its voltage-dependence. We delivered a depolarizing current step, in the presence of 1 µM TTX to block Na^+^-dependent APs, from a series of membrane potentials and measured the LTS. We found that the LTS was generated in FX relay cells, but only at a more hyperpolarized V_M_. While the LTS in WT relay cells was apparent at -60 mV, the LTS in FX relay cells only began to emerge at -70 mV (Fig. 4A-B; mixed-factor ANOVA, interaction between genotype and V_M_: F(2,20) = 8.958, p = 0.0017). Even when the LTS was evoked, the amplitude was significantly smaller in FX neurons compared to WT at both -60 mV (FX mean= 1.2 +/- 1.98 mV, WT mean= 13.7 +/- 7.32 mV) and - 70 mV (FX mean= 12.6 +/- 3.8 mV, WT mean= 19.3 +/- 3.4 mV) (Fig. 4B). The contribution of T-type Ca^2+^ channels to the LTS was confirmed by application of 50 µM Ni^2+^, a blocker of the T-type Ca^2+^ channels, to the extracellular saline (Fig. 4A)^34^.

**Figure 4:**
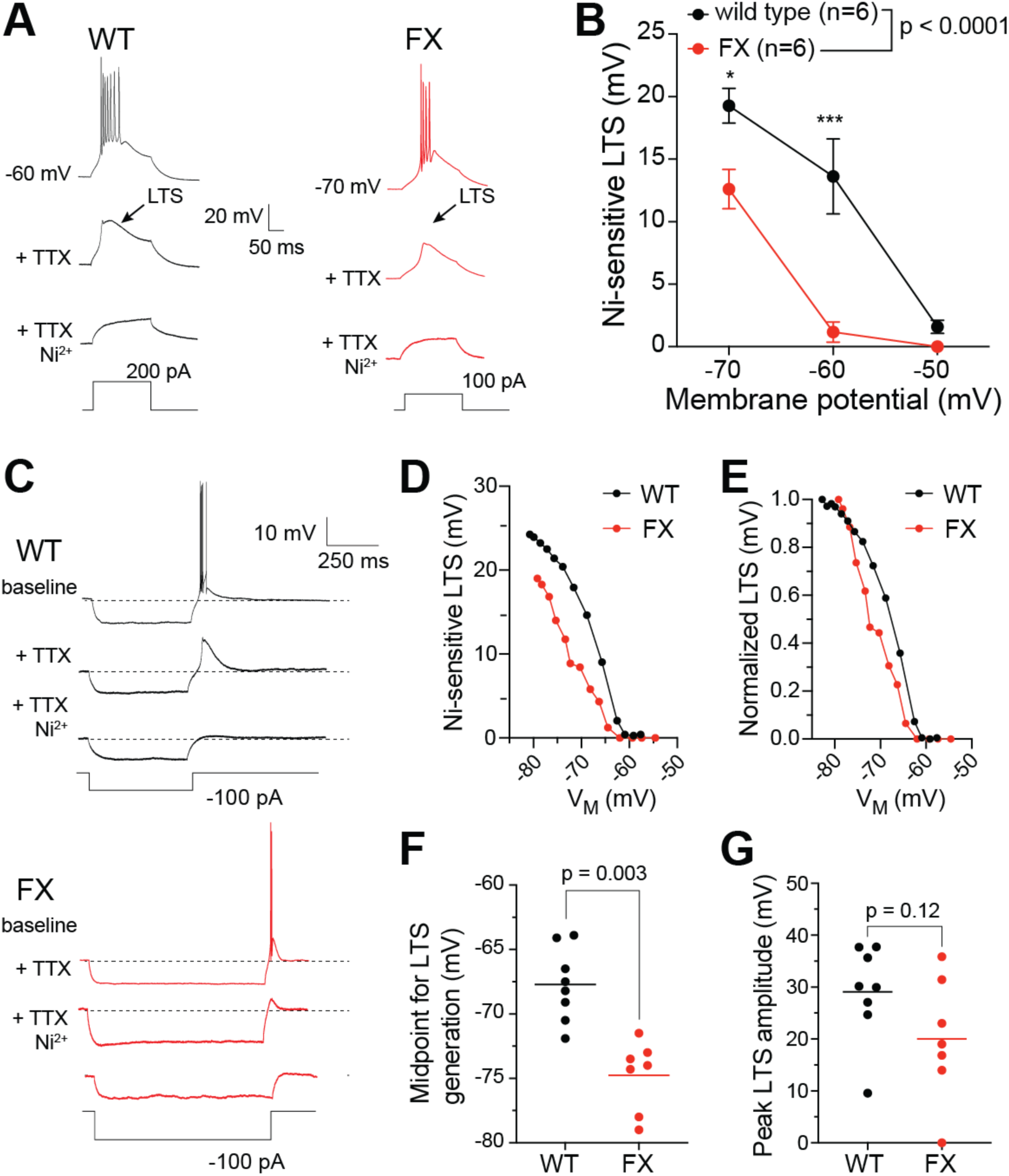
Low threshold spike generation is hyperpolarized in FX thalamic neurons. **A.** Representative traces showing a burst of AP, isolation of the LTS by TTX, and block of the LTS by Ni^2+^ **B.** Summary plot showing that LTS initiation is hyperpolarized in FX thalamic neurons. **C.** Example traces showing a rebound burst of APs following hyperpolarization, isolation of the LTS, and block of the LTS in WT and FX neurons. **D, E.** Individual WT and FX neurons showing raw (D) and normalized (E) LTS amplitude as a function of steady-state V_M_. **F**. Midpoint of LTS generation is hyperpolarized in FX compared to WT thalamic neurons. **G**, LTS amplitude between FX and WT thalamic neurons.

To examine in detail the altered voltage dependence of the LTS in FX mice, we measured the LTS that followed a series of hyperpolarizing current steps from a holding potential of -55 mV (Fig. 4C-E). In agreement with the experiment above, the voltage sensitivity of the rebound LTS was hyperpolarized in FX thalamic neurons compared to WT (Fig. 4F; unpaired t-test, t=4.880, df = 13, p = 0.0003). Similar to the LTS measured by depolarizing current steps (4a-b), the contribution of T-type Ca^2+^ channels to the LTS following hyperpolarization was confirmed by application of 50 µM Ni^2+^ (Fig. 4C)^34^.

## Discussion

We found that thalamic firing dynamics are profoundly disrupted in FX mice. *In vivo* recordings from awake mice revealed that while thalamic relay cells regularly transition between burst and tonic firing modes in the WT LGN, relay cells in FX LGN fire almost exclusively in the tonic mode. Furthermore, when FX relay cells do fire bursts, they contain fewer APs than those from WT relay cells. Using whole cell recordings in acute slices, we found that FX neurons continue to respond to depolarizing inputs in the tonic mode at membrane potentials which evoke bursts in WT neurons. FX neurons are capable generating burst APs, but only at more hyperpolarized membrane potentials. This change in firing pattern can be explained at least in part by a hyperpolarization of the voltage sensitivity of the LTS, which contributes to reduced burst frequency and duration in FX relay cells. These in vitro results provide a mechanistic basis for the dramatic reduction in burst frequency and duration FX LGN in vivo. While the LTS remains in FX relay cells, it requires greater membrane potential hyperpolarization for activation, which may only rarely occur *in vivo*. A smaller LTS when FX cells do transition to burst mode means less activation of voltage-dependent Na^+^ channels for driving APs, which will reduce the number of spikes in a burst.

There are additional mechanisms which may account for the functional disruption of bursts in FX LGN. Modulatory inputs to the LGN from brainstem cholinergic projections or thalamic reticular nucleus GABAergic projections could be altered in FX^12,35^. Properties of the retinal drive to the LGN which can evoke bursting could also be altered in FX mice^36^. These explanations are not mutually exclusive, as disruptions at both the network and cellular levels could contribute to the observed changes in thalamic firing dynamics in vivo. In addition, network-level disruptions in FX could explain why we observed lower mean firing rates in FX relay cells in vivo.

We used FX mice as a model for neurodevelopmental disruptions in ASD. FMR1 is conserved across species and its loss in humans causes a severe form of ASD, Fragile X syndrome. Recording in FX thalamus allowed us to probe the neural mechanisms contributing to the perceptual impairments commonly reported in ASD. We propose that a change in the pattern of firing modes in FX thalamus could disrupt sensory processing in neocortical regions by altering the normal patterns of thalamocortical relay. In the face of complex sensory environments, appropriate segmentation of task-relevant features from the background is essential to guide actions and avoid overwhelming the capacity of neocortical processing with irrelevant noise. What constitutes target and background in the sensory landscape can vary rapidly, making flexible control over what sensory information gets processed by neocortical circuits crucial for guiding optimal behaviors. A disruption of the neural mechanisms for dynamic gating of sensory relay could result in misallocation of neocortical processing resources to task-irrelevant stimuli, as well as failure to adequately detect, attend to, and interpret salient inputs. These predicted impairments are in line with perceptual deficits commonly seen in people with ASD. Across sensory modalities, ASD subjects display a generally reduced capacity to identify targets in noise or to direct attention to stimuli that have high salience for their neurotypical counterparts^2–7^.

Modulation of thalamic firing mode may control the selective gating of sensory information to cortex, as a function of both the “bottom-up” salience of peripheral inputs and “top-down” segmentation of task-relevant sensory information^14,23,25^. Experimental evidence supports the notion that the two firing modes play distinct roles in thalamocortical relay^13,19–21,25,26^. In addition, inhibitory inputs from the thalamic reticular nucleus to first order thalamic nuclei can alter the probability of bursting^12^. These findings implicate the dual firing modes as central to the dynamics of thalamocortical relay, with the capacity to be controlled by both the bottom-up salience of inputs and top-down, behaviorally relevant factors.

If the dynamics of thalamocortical relay do indeed play the hypothesized role in sensory processing, then rescuing normal thalamic firing patterns in FX, and potentially other forms of ASD, could allow people with ASD to better function in complex environments. Our results showing that a disruption in the voltage sensitivity of the LTS contributes to reduced burst frequency and duration in FX thalamus guides a potential therapeutic goal of rescuing normal T-type Ca^2+^ channel function and expression.

Previous in vitro studies demonstrated that rescuing FMRP expression in FX hippocampal and prefrontal neurons rescues HCN and potassium channel function^37,38^. Future work should explore whether rescuing FMRP expression in FX thalamic neurons restores the normal voltage-sensitivity of LTS generation. Ultimately, showing that reduced thalamic bursting contributes to perceptual impairments, and tying gain of function for voltage-sensitive ion channels to rescuing normal thalamic firing dynamics, could have therapeutic potential for people with ASD experiencing altered sensitivity to complex sensory environments.

## Methods

### *In vivo* extracellular recordings

All experiments were approved by the University of Texas at Austin’s Institutional Animal Care and Use Committee, which maintains Association for Assessment and Accreditation of Laboratory Animal Care accreditation. All methods were carried out in accordance with relevant guidelines and regulations.

### Animal preparation

Electrophysiology experiments were conducted using adult (P35 and older) C57/BL6J mice (WT) (Jackson Labs #000664) of both sexes (n = 11) and FMR1 knockout mice (FX) (Jackson Labs #003025) of both sexes (n=6), all housed on a 12-hour light-dark cycle.

For all surgical procedures, mice were anesthetized with isoflurane (2.5% induction, 1%–1.5% surgery) and given two preoperative subcutaneous injections of analgesia (5 mg/kg carprofen, 3.5 mg/kg Ethiqa) and anti-inflammatory agent (dexamethasone, 1%). Mice were kept warm with a water-regulated pump pad. Each mouse underwent two surgical procedures. The first was the mounting of a metal frame over the visual cortex using dental acrylic, used for head fixation of the mouse during experiments. The second was a 1-2mm diameter craniotomy centered 2.5mm posterior of bregma and 2mm lateral of midline in the right hemisphere followed by a durotomy. Exposed brain tissue was preserved prior to experiments with KWIK-SIL silicone adhesive (World Precision Instruments). Surgical procedures were always performed on a separate day from electrophysiological recordings.

### Electrophysiology

During all experiments, mice were awake, head restrained, and moved freely on a custom-built wheel. Mice were habituated to handling and head restraint over 1 week following recovery from the craniotomy in sessions increasing from 15 minutes to 1 hour. The axle of the wheel was connected to a quadrature encoder to track locomotion during the experiment.

LGN recordings were made using a Neuropixels 1.0 probe implanted acutely 2500 μm posterior of bregma and 2000 μm lateral of midline and lowered to 2500-3200 μm depth. The electrode tip was lowered to the ventral extent of the LGN as identified by clearly evoked visual responses in the neural activity. Raw traces were acquired at 30 kHz using the spikeGLX System.

Putative single units were isolated from the raw traces offline using Kilosort3^39^. We conducted further manual sorting of single units using the Phy GUI (https://github.com/cortex-lab/phy). All subsequent analysis of single unit spiking data was performed using custom MATLAB code.

Each spike from a single unit was defined as arising from the burst or tonic firing mode based on the pre and post ISI. The first AP in a burst is preceded by at least 100ms of silence, followed by APs separated by less than 4ms. A burst ends with an ISI greater than 4ms. In our data, as reported previously, the end of a burst tended to be followed by a period of at least 100ms of silence^29^. All spikes not meeting these criteria for bursting were classified as tonic spikes.

### Visual Stimuli

A monochrome LED projector by Texas Instruments (Keynote Photonics) with spectral peak at 525 nm was used to generate stimuli with a 60Hz refresh rate onto a Teflon screen which provides a near-Lambertian surface^40^. The screen was 12.5 cm high x 32 cm wide, equating to approximately 64° x 116° of visual angle. Stimuli were coded using the Psychophysics Toolbox extension in MATLAB.

Drifting gratings were presented at a spatial frequency of .02 cycles/degree, and a temporal frequency of 2 Hz, drifting at 0, 45, 90, 135 degrees, for 50 repeats of each stimulus. Each grating was preceded and followed by 1 second of a blank screen at uniform luminance. 50 repeats of 10 second natural movies were also presented with each repeat followed by a 6 second blank screen at uniform luminance. Movie 1 consisted of honeybees flying in a garden (Ian Nauhaus, UT Austin) and Movie 2 consisted of monkeys playing in snow (David Leopold, NIMH).

### *In vitro* whole-cell recordings

All experiments were approved by the University of Nevada, Las Vegas and the University of Texas at Austin Institutional Animal Care and Use Committees, which maintain Association for Assessment and Accreditation of Laboratory Animal Care accreditation. All methods were carried out in accordance with relevant guidelines and regulations.

### Acute slice preparation

Brain slices containing either the medial dorsal or visual thalamus were prepared and maintained as in prior studies^41,42^. Briefly, mice had free access to food and water and were housed in a reverse light-dark cycle of 12 hr on/ 12 hr off. Experiments used male 2–4-month-old wild type and *fmr1* knockout (FX) mice on a C57/Bl6 background (JAX: strain #000664). Mice were anesthetized using a ketamine/xylazine cocktail (100/10 mg/kg) and then underwent cardiac perfusions with ice-cold saline consisting of (in mM): 2.5 KCl, 1.25 NaH_2_PO_4_, 25 NaHCO_3_, 0.5 CaCl_2_, 7 MgCl_2_, 7 dextrose, 205 sucrose, 1.3 ascorbic acid, and 3 sodium pyruvate (bubbled constantly with 95% O_2_/5% CO_2_ to maintain pH at ∼7.4). The brain was removed and sliced into 300 µM coronal (for MD) or parasagittal (for LGN) sections using a vibrating tissue slicer (Vibratome 300, Vibratome Inc). The slices were placed in a chamber filled with artificial cerebral spinal fluid (aCSF) consisting of (in mM): 125 NaCl, 2.5 KCl, 1.25 NaH_2_PO_4_, 25 NaHCO_3_, 2 CaCl_2_, 2 MgCl_2_, 10 dextrose, 1.3 ascorbic acid and 3 sodium pyruvate (bubbled constantly with 95% O_2_/5% CO_2_) for 30 minutes at 35°C and then held at room temperature until time of recording.

### In vitro electrophysiology

Slices were placed individually, as needed, into a submerged recording chamber, continuously perfused with oxygenated extracellular saline containing (in mM): 125 NaCl, 3 KCl, 1.25 NaH_2_PO4, 25 NaHCO_3_, 2.0 CaCl_2_, 1.0 MgCl_2_, and 21 dextrose (pH 7.4) at 32-34°C. LGN and MD cells were viewed with a 60x water immersion objective and Dodt contrast. Ionotropic glutamatergic and GABAergic synaptic transmission were blocked with 20 µM DNQX, 25 µM D-AP5, and 2 µM gabazine. Current clamp recordings were made using a Dagan BVC-700 amplifier and acquisition software Axograph X (Axograph) or SutterPatch (Sutter). Data were sampled at 20-50 kHz, filtered at 3 kHz, and then digitized by either an InstruTECH ITC-18 interface (HEKA) or Dendrite interface (Sutter). The internal recording solution consisted of (in mM): 135 K-gluconate, 10 HEPES, 7 NaCl, 7 K_2_-phosphocreatine, 0.3 Na-GTP, 4 Mg-ATP (pH corrected to 7.3 with KOH). Recording pipettes had a resistance of 4-6 MΩ when filled with the pipette solution. During current clamp recordings, series resistance was monitored throughout, and the experiment was discarded if it exceeded 30 MΩ or varied by >20%. The LTS was isolated by blocking Na^+^-dependent APs with TTX (1 µM) during whole-cell recordings. Following LTS recordings, NiCl_2_ (50 µM) was applied to confirm that the measured LTS was mediated by T-type Ca^2+^channels^34^.

---

**Supplementary Table 1:** Neuronal Properties.

|  | wild type | Fmr1 KO | p-value | n |
| --- | --- | --- | --- | --- |
| ***Passive Properties*** |  |  |  |  |
| Resting V<sub>M</sub> (mV) | -59.3 ± 1.89 | -58.6 ± 2.39 | 0.8262 | 11, 9 |
| Input resistance (MΩ) | 111.3 ± 14.49 | 129.1 ± 15.16 | 0.2541 | 11, 9 |
| $\tau_M$ (ms) | 21.1 ± 3.62 | 27.7 ± 3.42 | 0.232 | 6, 6 |
| Voltage sag (%) | 13.6 ± 3.66 | 13 ± 4.52 | 0.92 | 7, 6 |
| ***Active Properties*** |  |  |  |  |
| AP threshold (mV) | -38.5 ± 2.77 | -37.4 ± 3.19 | 0.8003 | 8, 6 |
| AP amplitude (mV) | 81.8 ± 3.35 | 81.5 ± 4.53 | 0.9414 | 8, 6 |
| AP width (μs) | 698 ± 23.4 | 771 ± 65.8 | 0.2606 | 8, 6 |
| AP maximum rate of rise (mV/ms) | 269 ± 16.4 | 278 ± 33.2 | 0.8099 | 8, 6 |
| AP maximum rate of fall (mV/ms) | -122 ± 5.9 | -107 ± 9.8 | 0.1817 | 8, 6 |

## Acknowledgements

The authors wish to thank Carrie Barr for animal care and laboratory management, and Laura Colgin for helpful feedback on the manuscript.

Research funding was provided by NIH-R01-EY028657 (N.J.P.), NIH-R01-MH131317 (D.H.B.), and NIH-5-T32-EY-021462-12 (R.T.O.).

